# Neural Speech-Tracking During Selective Attention: A Spatially Realistic Audiovisual Study

**DOI:** 10.1101/2023.12.27.573349

**Authors:** Paz Har-shai Yahav, Eshed Rabinovitch, Adi Korisky, Renana Vaknin Harel, Martin Bliechner, Elana Zion Golumbic

## Abstract

Paying attention to a target talker in multi-talker scenarios is associated with its more accurate neural-tracking relative to competing non-target speech. This “neural-bias” to target speech has largely been demonstrated in experimental setups where target and non-target speech are acoustically controlled and interchangeable. However, in real-life situations this is rarely the case. For example, listeners often look at the talker they are paying attention to while non-target speech is heard (but not seen) from peripheral locations. To enhance the ecological-relevance of attention research, here we studied whether neural-bias towards target speech is observed in a spatially realistic audiovisual context, and how this is affected by switching the identity of the target talker.

Group-level results show robust neural-bias towards target speech, an effect that persisted and generalized after switching the identity of the target talker. In line with previous studies, this supports the utility of the speech-tracking approach for studying speech processing and attention in spatially realistic settings. However, a more nuanced picture emerges when inspecting data of individual participants. Although reliable neural speech-tracking could be established in most participants, this was not correlated with neural-bias or with behavioral performance, and >50% of participant showed similarly robust neural tracking of both target and non-target speech.

These results indicate that neural-bias toward the target is not a ubiquitous, or necessary, marker of selective attention (at least as measured from scalp-EEG), and suggest that individuals diverge in their internal prioritization among concurrent speech, perhaps reflecting different listening strategies or capabilities under realistic conditions.

**Significance Statement:** This work contributes to ongoing efforts to study the neural mechanisms involved in selective attention to speech under ecologically relevant conditions, emulating the type of speech materials, multisensory experience, and spatial realism of natural environments. Group-level results show that under these more realistic conditions, the hallmark signature of selective attention – namely the modulation of sensory representation, and its robustness to switches in target-identity – is conserved, at least at the group level. At the same time, results point to an underlying diversity among participants in how that this modulation manifests, raising the possibility that differences in listening strategies, motivation or personal traits lead to differences in the way that individuals encode and process competing stimuli, under ecological conditions.

## Introduction

Effectively directing attention to a particular talker, and prioritizing its processing over competing non-target speech, can be challenging. Selective attention to speech to has been associated at the neural level with enhanced neural-tracking of target speech, compared to non-target speech. This “neural-bias” for target speech has been demonstrated in numerous EEG and MEG studies of selective attention to speech (Ding et al., 2012; Fiedler et al., 2019; Kerlin et al., 2010; J. A. O’Sullivan et al., 2015; Zion Golumbic, Ding, et al., 2013), and mirrors similar effects of selective attention in modulating sensory responses to simpler stimuli (Broadbent, 1958; Hillyard et al., 1973; Näätänen et al., 1992; Treisman, 1969; Tsotsos et al., 1995). However, to date, this effect has been mostly studied under conditions that don’t fully capture the real-life challenge of attention to speech. For example, the speech materials used in many studies are often comprised either of short, context-less utterances (Brungart, 2001; Humes et al., 2017) or recordings of audiobooks that are highly edited and professionally rehearsed and recorded (Fiedler et al., 2019; Fu et al., 2019a), functioning more as “spoken texts” than as natural speech. In contrast, natural speech is continuous, contextual and is produced on-the-fly resulting in added disfluencies, pauses and repetitions (Agmon et al., 2023).

Another non-ecological aspect of many studies is that speech is presented only auditorily, often in a dichotic manner where the audio from different talkers is presented to different ears (Aydelott et al., 2015; Bentin et al., 1995; Brodbeck et al., 2020; Cherry, 1953; Kaufman & Zion Golumbic, 2023; Makov et al., 2022). However, in many real-life situations, listeners also look at the talker that they are paying attention to, hence target speech is often audiovisual by nature and is emitted from a central location relative to the listener. In contrast, other non-target talkers are – by default – heard but not seen (unless listeners overtly move their head/eyes), and their audio emanates from peripheral spatial locations. Accordingly, under spatially realistic audiovisual conditions, there are stark qualitative differences between the sensory features of target and non-target speech, which likely assist listeners in focusing their attention appropriately (Fleming et al., 2021). Supporting this, it has been shown that having corresponding visual input of a talker facilitates speech processing as well as selective attention (Ahmed, Nidiffer, & Lalor, 2023; Ahmed, Nidiffer, O’Sullivan, et al., 2023; Grant & Seitz, 2000; Haider et al., 2024; Karthik et al., 2024; Schwartz et al., 2004; Sumby & Pollack, 1954; Wikman et al., 2024) and also improves the precision of neural speech-tracking (Crosse et al., 2015; Fu et al., 2019b; Zion Golumbic, Cogan, et al., 2013).

The current study is part of ongoing efforts to increase the ecological validity of selective attention research and to advance our understanding of how the brain processes and prioritizes competing speech in the type of circumstances encountered in real-life (Brown et al., 2023; Freyman et al., 2001; Keidser et al., 2020; Ross et al., 2007; Shavit-Cohen & Zion Golumbic, 2019; Tye-Mmurray et al., 2016; Uhrig et al., 2022). We capitalize on the potential of the speech-tracking approach for gaining insight into how the brain encodes and represents concurrent, continuous and natural speech stimuli (Ding et al., 2012; Kaufman & Zion Golumbic, 2023; Mesgarani & Chang, 2012; Zion Golumbic, Ding, et al., 2013). To our knowledge, only a handful of previous studies have measured neural speech-tracking in a spatially-real audiovisual selective attention paradigm (A. E. suppullivan et al., 2019; Wang et al., 2023). In one such study, O’Sullivan et al. (2019) found that it is possible to determine from the neural signal whether a listener is paying attention to the talker they are looking at or if they are ‘eavesdropping’ and paying attention to a peripheral talker whom they cannot see. These results nicely demonstrate the dissimilarity of the neural representation of audiovisual target speech and concurrent audio-only non-target speech. However, they leave open the question of the degree to which the brain suppresses irrelevant speech and exhibits “neural-bias” for preferential encoding of target speech under spatially realistic audiovisual conditions.

There is a long-standing theoretical debate about how selective attention affects the neural representation of non-target stimuli. One possibility is that non-target speech is attenuated at an early sensory level structuring (Broadbent, 1958; Carlyon, 2004; Ding et al., 2018; Treisman, 1960), and the degree to which non-target speech is represented/attenuated is thought to reflect the efficacy of selective attention. Alternatively, target and non-target speech can be co-represented at the sensory level, with selection occurring only at later stages (e.g., the level of linguistic/semantic processing; late-selection (Deutsch & Deutsch, 1963; Murphy et al., 2013). As noted, numerous studies have demonstrated reliable “neural-bias” in the sensory representation of concurrent speech in dichotic listening paradigms, showing that the acoustic envelope of target speech is tracked more precisely than non-target speech (Fiedler et al., 2019; Fuglsang et al., 2017; Har-shai Yahav et al., 2023; Kerlin et al., 2010; J. A. O’Sullivan et al., 2015; Zion Golumbic, Ding, et al., 2013). However, although this effect is robust when averaging across multiple participants, recently, Kaufman & Zion Golumbic (2023) showed that this effect is driven by ∼30% of participants, whereas the majority of participants do not show reliable neural-bias, but exhibit comparable neural representation for both target and non-target speech in bilateral auditory cortex. This raised the possibility that suppression of non-target speech, at least at the sensory level, may not be a necessary component of selective attention, and that differences between individuals may reflect different listening strategies or capability for multiplexed listening. However, that study, like most previous work in the field, used an auditory-only dichotic listening design, which does not capture the spatial realism of real-life audiovisual contexts.

Here sought to replicate and extend our previous work using a more ecologically valid spatially-realistic audiovisual design, and to study the relative representation of target and non-target speech in the brain under these circumstances. We simulated a common real-life situation in which individuals pay attention to a talker whom they can see (in this case, watching a video-recording of a lecture), but also hear another talker off to the side, who they cannot see and are asked to ignore. We presented the audio of both target and non-target in a free-field fashion, from their respective spatial locations, rather than through earphones, to ensure realistic spatial propagation of the sound. Importantly, we used unedited recordings of actual lectures delivered by academics for the general public, to preserve the natural properties of the speech. In addition, mid-way along the experiment we switched between the target and non-target talkers, to study how this affected listeners and to test the generalizability of results across talkers and over time.

We recorded participants neural activity using electroencephalography (EEG) and analyzed their neural-tracking of target and non-target speech, focusing both on group-averages, as is common in the field, as well as on individual-level data (Ding et al., 2012; Ding & Simon, 2012; Fuglsang et al., 2017; Kaufman & Zion Golumbic, 2023; Rosenkranz et al., 2021b). We use data-driven statistics to determine the degree to which each talker is represented in the neural signal as well as the “neural-bias” towards target speech. We also tested the reliability of results between the first and second half of the experiment, and tested if switching the identity of the target talker affected the pattern of neural-tracking.

## Materials and Methods

### Participants

We collected data from 24 adult volunteers (16 female, 8 male), ranging in age between 19 and 34 (M=23.83, SD=±3.42). All participants were fluent Hebrew speakers with self-reported normal hearing and no history of psychiatric or neurological disorders. The study was approved by the IRB ethics board of Bar-Ilan University, and participants gave their written informed consent prior to the experiment. Participants were either paid or received course credit for participating in the experiment. Data from one participant was excluded from all analysis due to technical issues during EEG recording, therefore all further analyses are reported for N=23.

### Speech stimuli

The stimuli consisted of two 20-minutes video recordings of a public lecture on popular science topics, one delivered by a male talker and the other by a female talker. Each video recording included the lecturer as well as the slides accompanying the talk. Both talkers gave their approval to use these materials for research purposes. Lecture videos were segmented, edited and volume equated using Matlab (The MathWorks), Filmora software (filmora.wondershare.net), Audacity software (version 3.2.1, www.audacityteam.org) and FFMPEG (www.ffmpeg.org). Lectures were cut into 63 segments ranging between 22 to 40 seconds each (lecture 1: M=31.83, SD=±4.2, lecture 2: M=30.68, SD=2.24). The varied lengths were necessary to ensure that the segments did not cut-off the lecture mid-sentence or mid-thought. Ultimately, the experiment included only 42 segments from each lecture, with 21 un-used segments from the middle of the lecture (see experimental procedure).

### Experimental procedure

The experiment was programmed and presented to participants using the software OpenSesame (Mathôt et al., 2012). Participants were seated on a comfortable chair in a sound attenuated booth and were instructed to keep as still as possible. Participants viewed a video lecture (target), presented segment-by-segment, on a computer monitor in front of them with the lecture audio presented through a loudspeaker placed behind the monitor. They were instructed to pay attention to the video and after every three segments, were asked three multiple-choice comprehension questions, regarding the content of the recent segments (one question per segment). Participants received feedback regarding the correctness of their responses, and indicated via button press when they were ready to continue to the next segment. In addition to the target lecture, audio from an additional non-target lecture was presented through a loudspeaker placed on their left side. Segments of non-target speech began 3 seconds after the onset of the target speech, and included a volume ramp-up of 2 seconds (Figure 1B). Both loudspeakers were placed at the same distance from participants’ head (approximately 95cm). We chose to present non-target speech only from the left side, rather than counterbalanced across both sides, to ensure sufficient amount of data for TRF estimation, without doubling the experiment length.

**Figure 1.**
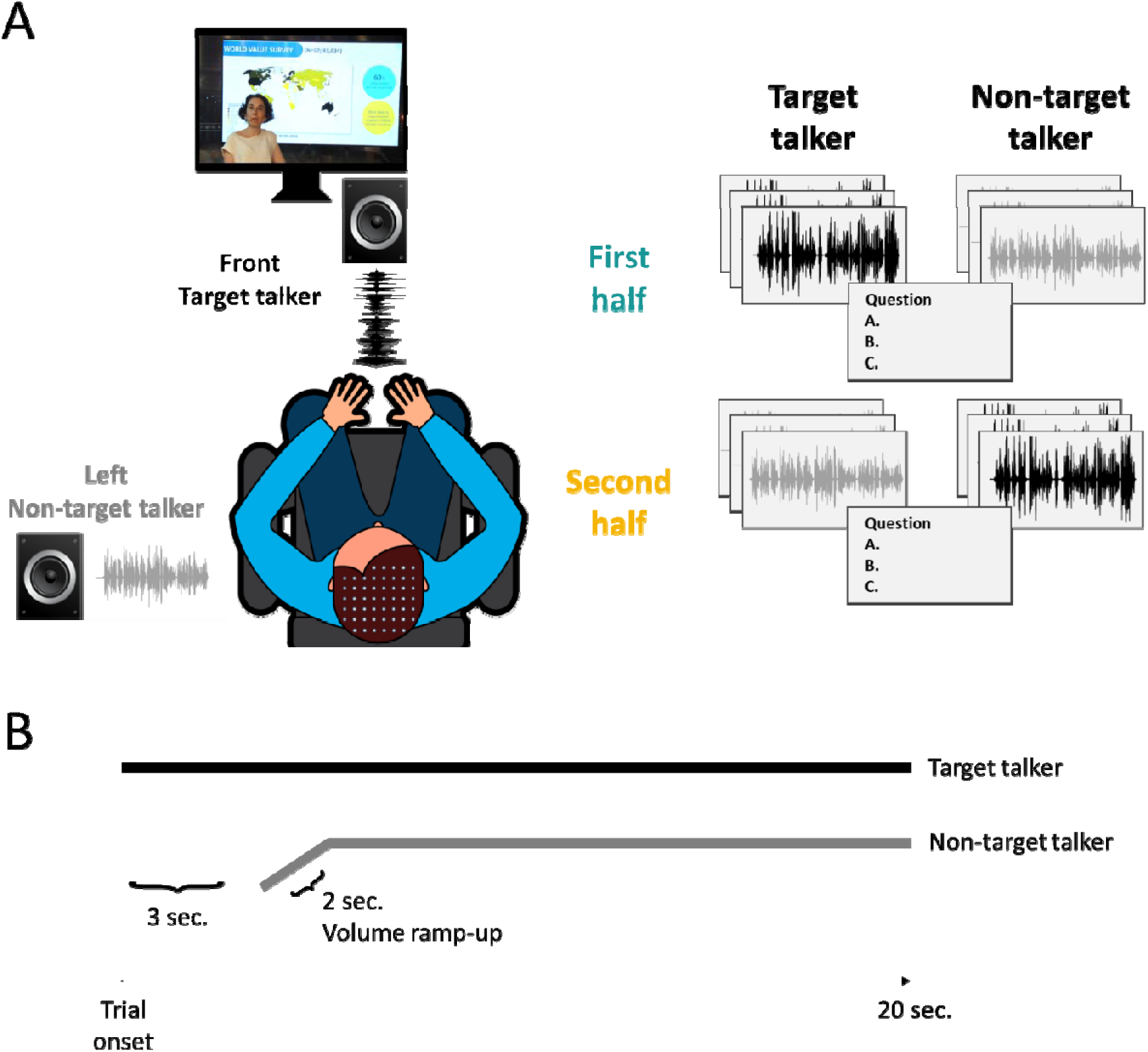
Experimental setup. A. Two lectures were presented simultaneously, with one lecture (target talker) displayed on the screen and audio emitted through the front loudspeaker. The other lecture (non-target talker) was played audio-only through the left loudspeaker. Participants were instructed to focus their attention on the lecture presented on the screen. Critically, in the middle of the experiment, the stimuli switched such that the lecture that was played from the side and had been non-target in the first half, was presented as a video in the second half and became the target talker, whereas the target talker from the first half was presented in the second half from a loudspeaker on the side and became non-target. Participants answered comprehension questions regarding the target lecture after every three trials. B. Single trial illustration. Target speech began at trial onset, non-target speech began 3 seconds after onset and included a 2-second volume ramp-up.

In the middle of the experiment, the stimuli were switched: The lecture that had been the non-target became the target lecture and was presented as a video in the second half, whereas the lecture that had been the target, was presented as audio-only from a loudspeaker on the left, and was the non-target (Figure 1A). Importantly, different portions of each lecture were presented in each half of the experiment. When a lecture was designated as the target, it started from the beginning to ensure optimal comprehension, and continued for 21 consecutive segments. When a lecture was designated as the non-target (and presented only auditorily), the last 21 segments of the lecture were played, also in consecutive order. In this way, segments from each lecture served both as target- and non-target speech in different parts of the experiment (thus sharing the talker-specific attributes and general topic), but none of the content was repeated. The order of the starting lecture (male/female talker) was counterbalanced across participants, and participants were not informed in advance about the switch. Audio from on-ear microphones and eye movements were also recorded during the experiment, but their analysis is outside the scope of this study.

### EEG data acquisition

Electroencephalography (EEG) was recorded using a 64 Active-Two system (BioSemi; sampling rate: 2048 Hz) with Ag-AgCl electrodes, placed according to the 10–20 system. Two external electrodes were placed on the mastoids and served as reference channels. Electrooculographic (EOG) signals were simultaneously measured by 4 additional electrodes, located above and beneath the right eye and on the external side of both eyes.

### Behavioral data analysis

Behavioral data consisted of accuracy on the comprehension questions asked about each segment. These values were averaged across trials, separately for each half of the experiment, and for each participant. We used a two-tailed paired t-test to evaluate whether accuracy rates differed significantly before and after the talker-switch (first and second half).

### EEG preprocessing and speech-tracking

EEG preprocessing and analysis were performed using the Matlab-based FieldTrip toolbox (Oostenveld et al., 2011), as well as custom written scripts. Raw data was first re-referenced to the linked left and right mastoids and was bandpass filtered between 0.5-40Hz (4^th^ order zero-phase Butterworth filter). Data were then visually inspected and gross artifacts (that were not eye-movements) were removed. Independent component analysis (ICA) was performed to identify and remove components associated with horizontal or vertical eye-movements as well as heartbeats (identified through visual inspection). Any remaining noisy electrodes that exhibited either extreme high-frequency activity or low-frequency drifts, were replaced with the average of their neighbors using an interpolation procedure (ft_channelrepair function in the Fieldtrip toolbox). The clean data was then cut into trials, corresponding to portions of the experiment in which a single segment of the lecture was presented. These were divided according to which half of the experiment there were from – before and after switching the target talker (first and second half).

To estimate neural responses to the speech from the two simultaneous lectures we performed speech-tracking analysis, using both an encoding and a decoding approach. We estimated multivariate linear Temporal Response Functions (TRFs) using the mTRF MATLAB toolbox (Crosse et al., 2016), which constitutes a linear transfer function describing the relationship between a particular feature of the stimuli (S) and the neural response (R) recorded when hearing it. Here S was a matric comprised of the broadband envelopes of the two speech stimuli presented in each trial, and they were treated as separate regressors in a multivariate regression model. Envelopes were extracted using an equally spaced filterbank between 100-10,000 Hz, with 20 frequency bands, based on Liberman’s cochlear frequency map (Liberman, 1982). The narrowband filtered signals were summed across bands after taking the absolute value of the Hilbert transform for each one, resulting in a broadband envelope signal. The R used here was the continuous cleaned EEG data, bandpass filtered again between 1 and 20 Hz (4^th^ order zero-phase Butterworth filter), since the speech-tracking response consists mostly of low-frequency modulations. S and R were aligned in time and down-sampled to 100 Hz for computational efficiency. The 5 first seconds of each trial were excluded from data analysis, due to the differences in onset and ramp-up period (Figure 1B). In addition, the first trial and the trial immediately after the talker-switch were omitted from data analysis, to avoid confounding effects associated with attentional ambiguity and adjustment to the new target talker. This resulted in a total of 40 trials analyzed (20 per half).

Univariate and multivariate encoding and decoding models were optimized separately for each half of the experiment. In the encoding approach, linear TRFs are estimated reflecting the neural response at each electrode for each of the two simultaneously presented stimuli, and the predictive power of the model reflects how it predicts the actual neural response recorded. In the decoding approach, the neural data is used to reconstruct the envelope of each speech-stimulus. TRF predictive power values (encoding) and reconstruction accuracies (decoding) were assessed using a leave-one-out cross validation protocol. In each iteration, all trials except one were randomly selected to train the model (train set), which was then used to predict either the neural response at each electrode (encoding) or the two speech envelopes (decoding) in the left-out trial (test set). The predictive power of the encoding model is the Pearson’s correlation (r-value) between the actual neural response in the left-out-trial and the response predicted by the model. The decoding reconstruction accuracies is calculated separately for the two speech stimuli presented in the left-out-trial (target and non-target speech) and is the Pearson’s correlation (r-value) between the reconstructed envelope of each and the actual envelope. The reported TRFs, predictive power values and reconstruction accuracies are the averages across all iterations.

Encoding TRFs were calculated over time lags ranging from −150 (pre-stimulus) to 450 ms, and the decoding analysis used time lags of 0 to 400 ms. To prevent overfitting of the model, a ridge parameter was chosen as part of the cross-validation process (λ -predictive power). This parameter significantly influences the shape and amplitude of the TRF and therefore, rather than choosing a different λ for each participant (which would limit group-level analyses), a common λ value was selected for all participants, which yielded the highest average predictive power, across all channels and participants (see also Har-shai Yahav et al., 2023; Kaufman & Zion Golumbic, 2023). For both the encoding and decoding models this optimal value was λ=1000 a. Note that decoding results were highly similar for λ’s that were optimized separately for each participant (data not shown).

### EEG Statistical analysis Group level-statistics

The statistical significance of the predictive power and reconstruction accuracy of the encoding and decoding models were evaluated using permutation tests. For this, we repeated the encoding/decoding analysis procedure on shuffled S-R data where speech-envelopes presented in one trial (S) were paired with the neural response recorded in a different trial (mismatched R). This procedure was repeated 100 times, yielding a null-distribution of predictive power/reconstruction accuracy values that could be obtained by chance. The real data was then compared to this null-distribution and if it fell within the top 5% was considered statistically significant. To compare the speech-tracking response across conditions, we conducted a 2x2 ANOVA a Bayesian factor analysis with repeated-measures to compare the predictive-power and reconstruction accuracies obtained for target vs. non-target speech, in each half of the experiment (before vs. after the target talker switch) (JASP-Team, 2022; version 0.16.3) (priors distribution parameters: Uniform Cauchy distribution with a scale parameter of r = 0.5, random effects r = 1, scale covariates r=0.354) To test the generalizability of the speech-tracking patterns between the two halves of the experiment, we also tested how well decoders that were trained on data from one half of the experiment (either on target or non-target speech) could be used to accurately predict the stimuli presented in the other half of the experiment based on the neural data.

### Individual level-statistics

Statistical analysis of individual-level data focused only on reconstruction accuracies (decoding approach), since this approach integrates responses across all electrodes, yielding a simpler metric for statistical analysis and avoiding multiple comparisons. We conducted a series of permutation tests to obtain data-driven statistics in each individual participant, designed to address different questions, as illustrated in Figure 2.

**Figure 2.**
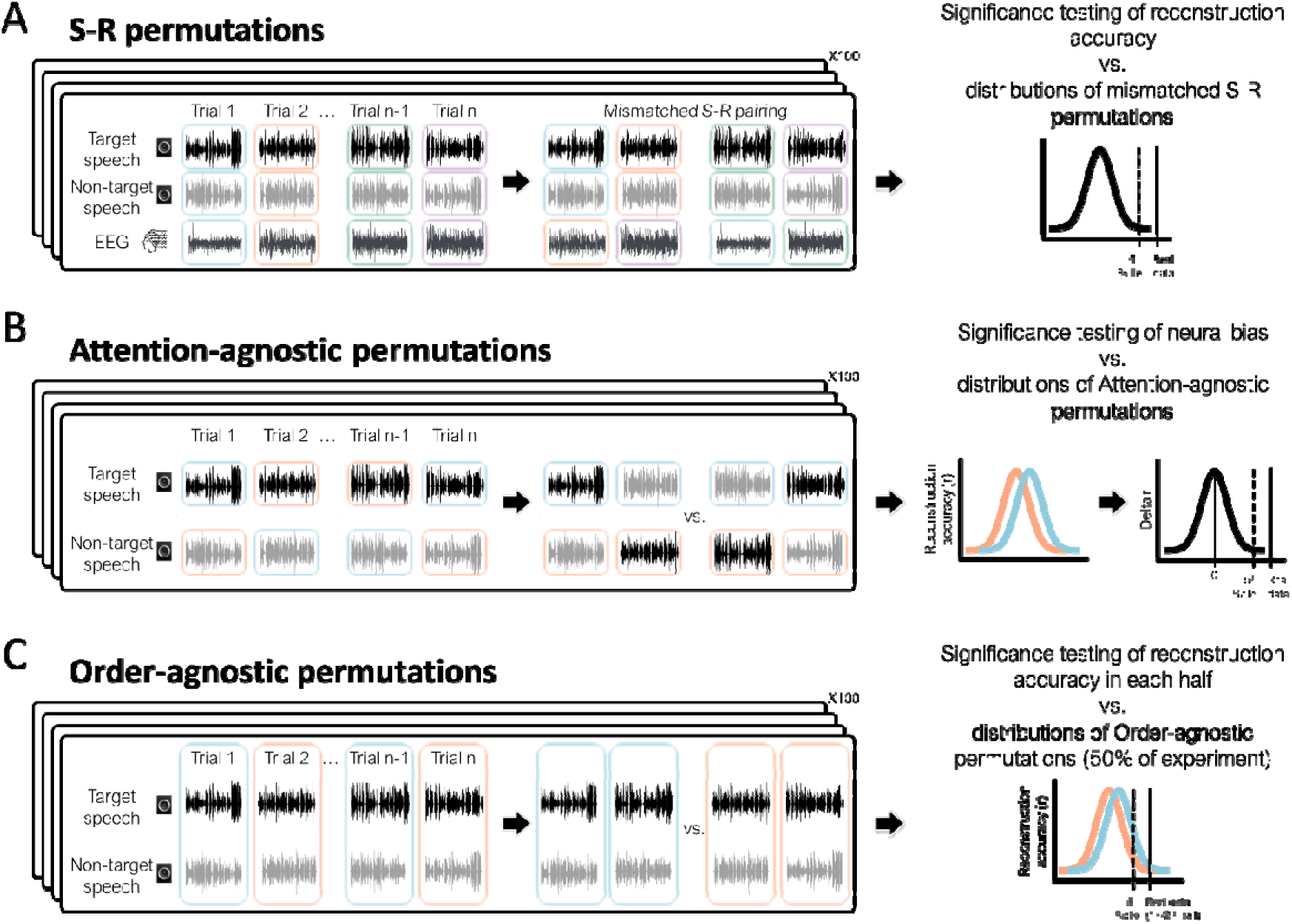
Data-driven permutation tests for individual-level statistics. Three permutation tests were designed to assess statistical significance of different results in individual participants. The black rectangles in all panels show the original data organization on the left and the re-labeling for permutations test on the right. **A. S-R permutation test.** In each permutation, the pairing between acoustic envelopes (S) and neural data responses (R) was shuffled across trials such that speech-envelopes presented in one trial (both target & non-target speech) were paired with the neural response (R) from in a different trial. This yields a null-distribution of reconstruction accuracies that could be obtained by chance, to which the real-data can be compared (right). **B. Attention-agnostic permutation test.** In each permutation, the target and non-target speech stimuli were randomly re-labeled to create attention-agnostic regressors that contain 50% target speech and 50% non-target speech. The reconstruction accuracy for each regressor was estimated and the difference between them is used to create a null-distribution to which the neural-bias index can be compared (right). **C. Order-agnostic permutation test.** In each permutation, trials were randomly re-labeled and separated into two order-agnostic groups consisting of 50% trials from the first half of the experiment and 50% trials from the second half. The reconstruction accuracy for each group of trials was estimated and the difference between them is used to create a null-distribution to which the real data from each half of the experiment can be compared (right).

First, we assessed whether the speech reconstruction accuracies obtained for both target and non-target speech were significantly better than those that could be obtained chance. To this end, we used S-R permutations, similar to those used for group-level statistics, in which we shuffled the pairing between acoustic envelopes (S) and neural data responses (R) (Figure 2A). Reconstruction values were assessed in 100 permutations of mismatched S-R combinations, yielding a null-distribution from which we derived a personalized chance-level value for target and non-target speech, for each participant (the top 5 of the null-distribution).

Second, we assessed whether the <u>difference</u> in the reconstruction accuracy for the target and non-target talker could reliably be attributed to their task-relevance (referred to as the “Neural-Bias index”). For this, we followed the procedure introduced by Kaufman & Zion Golumbic (2023) to create an “attention-agnostic” null-distribution of neural-bias indices (Figure 2B). Specifically, for each participant we created 100 permutation in which the two speech stimuli were randomly re-labeled so that the stimuli represented in each regressor was 50% target and 50% non-target. Multivariate decoders were trained on this re-labeled data and reconstruction accuracy values were estimated for each regressor, and difference between them was used to create an attention-agnostic distribution of differences between reconstruction accuracies. The real neural-bias index for each participant was normalized (z-scored) relative to this null-distribution, and participants with a z-score > 1.64 were considered as exhibiting a significance neural-bias towards the target speech (p<0.05 one tailed). We chose this normalization continuous approach, rather than using a cutoff value, which allows us to present the distribution of neural-bias values across participants. We note that the approach used here to assess differences in speech-tracking of target and non-target here differs from the auditory attention-decoding (AAD) approach used in similar studies to identify which of two speech stimuli belongs to the target talker (Fuglsang et al., 2017; Mirkovic et al., 2015; A. E. O’Sullivan et al., 2019; J. A. O’Sullivan et al., 2015; Teoh & Lalor, 2019). In those studies, a decoder <u>trained only on target speech</u> is used to predict the envelope of non-target speech (and vice versa), which assesses the similarity/differences between the decoders estimated for each stimulus. However, this approach is less appropriate in the current study, where we were interested in assessing how accurately each speech stimulus is represented in the neural signal, even if the spatio-temporal features of their decoders are different (see Extended Data Figure S1 for a direct comparison of these approaches and discussion of their utility for different purposes).

The two analyses described above (Figure 2A&B) were performed using data from the entire experiment, as well as on data from each half of the experiment separately. We then performed a third permutation test to assess whether speech reconstruction accuracies and the Neural-Bias index differed significantly in the two halves of the experiment, i.e., before vs. after the talker-switch. For this, we conducted an “order-agnostic” permutation test where trials were randomly re-labeled so that the data included in each regressor were 50% from the first half of the experiment and 50% from the second half (Figure 2C). Multivariate decoders were trained on this re-labeled data and reconstruction accuracy values were estimated for each regressor and this procedure was repeated

100 times, yielding a null-distribution. A participant was considered as showing a significant difference in neural-tracking of either the target or non-target talker, or different in neural-bias, if the their real-data fell in the top 5^%tile^ of the relevant null-distribution.

## Results

### Behavioral data

Accuracy on answering comprehension questions about the content of the lecture was significantly above chance [M=0.845, SD=±0.076; t(22)=32.178, p<10]. No significant differences in performance were observed between the first half (M=0.846, SD=±0.09) and second half (M=0.856, SD=±0.1) of the experiment [t(22)=-0.42, p=0.68], as shown in Figure 3.

**Figure 3.**
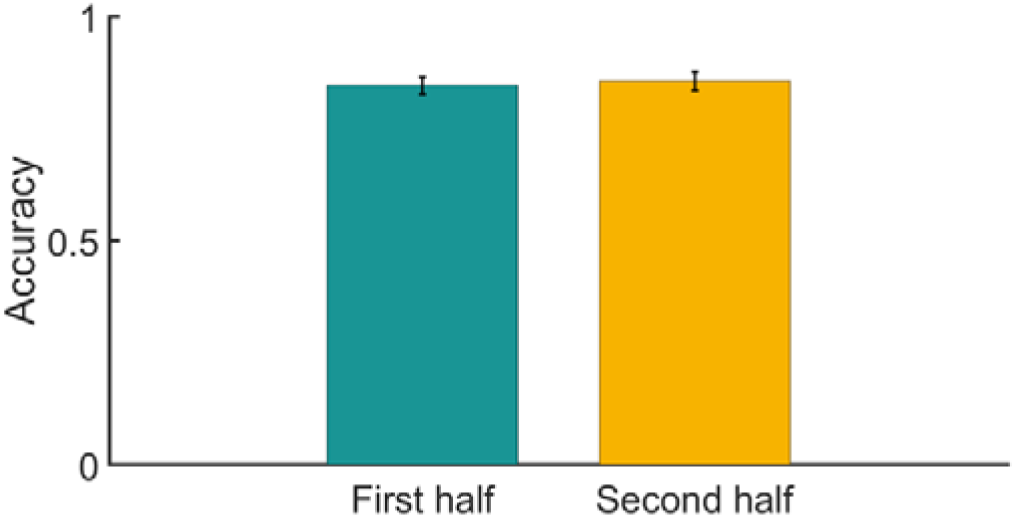
Behavioral results. Averaged accuracy rates across trials and participants, for multiple-choice comprehension questions, separately for the first (green) and second (yellow) half of the experiment. Error-bars denote SEM across participants.

### Speech-tracking analysis

#### Group-level results

Figure 4 shows the results of the encoding approach. TRF models were estimated for target and non-target speech, trained on data from the entire experiment and also separately on each half of the experiment (before and after talker-switch). As expected, the predictive power followed the common centro-frontal topography characteristic of auditory responses in EEG (Figure 4A), and was significant compared to a null-distribution (p<0.01). The TRF for target speech showed three prominent peaks – at 60ms, 110ms and 170ms – which is in line with previous studies and are thought to reflect a cascade of information flow from primary auditory cortex to associative and highly-order regions (Brodbeck, Presacco, et al., 2018). The TRF for non-target speech was also robust, however showed only a single positive peak, around 70 ms, which likely reflects its early sensory encoding, but not the two later peaks. Although the TRFs estimated for target and non-target speech are not directly comparable (due to the many sensory differences between them), they did differ in the amount of variance they explained in the neural signal [predictive power of univariate models for target vs. non-target, averaged across all electrodes; F(22)=11.5, p=0.003] and including both regressors in a multivariate model explained significantly more of the variance in the neural signal than either univariate model alone (t-test between average predictive power: multivariate vs. target only: t=2.97, p=0.007; multivariate vs. non-target only: t=5.9, p<10⁻⁵; Figure 4B).

**Figure 4.**
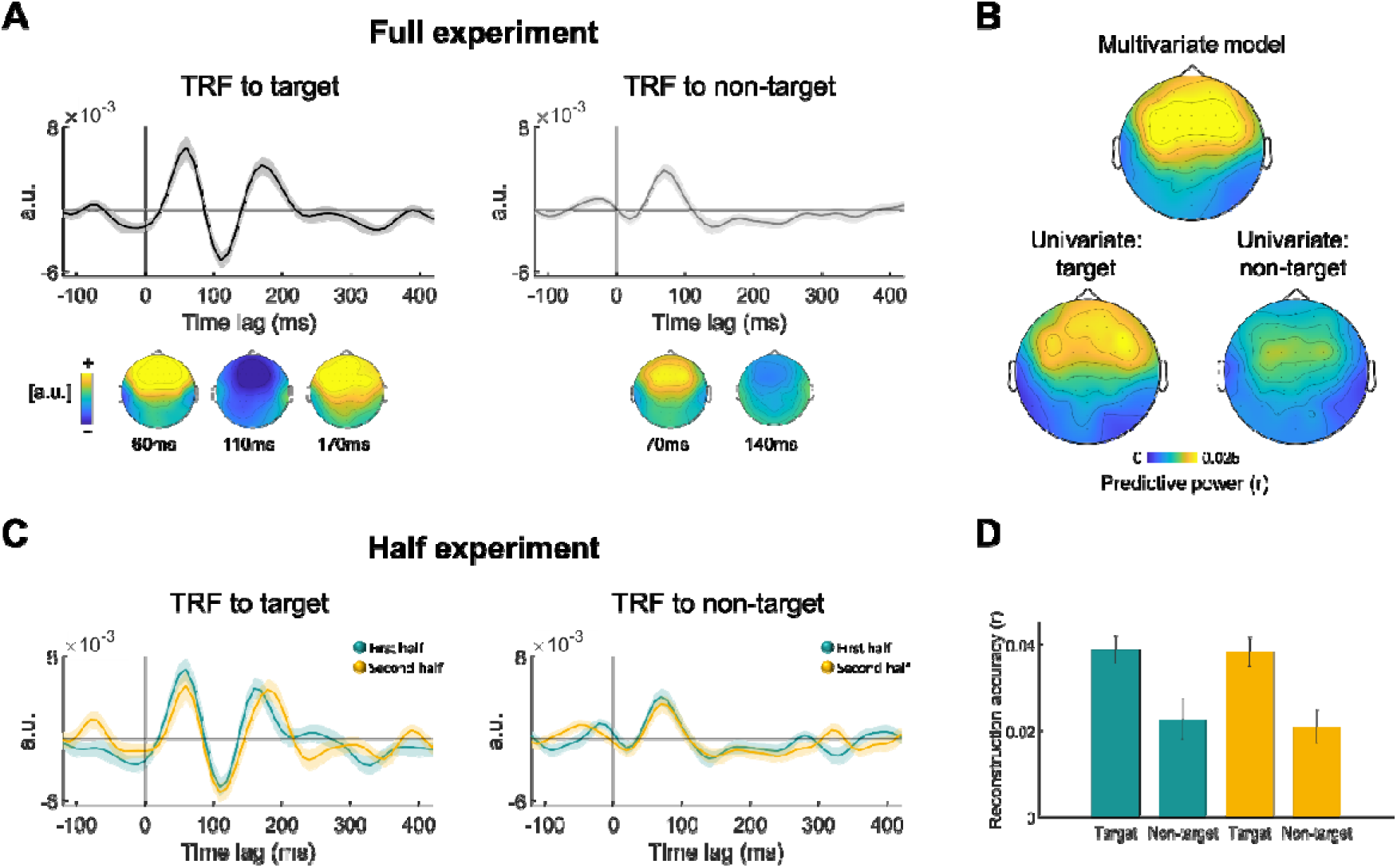
Neural bias: Group-level results. A. TRF encoding models across all experimental trials, plotted from electrode ‘Fz’, separately for target and non-target speech. Shaded highlights denote SEM across participants (top). Topographic distribution of the TRF main peaks, plotted separately for target and non-target speech (bottom). B. Topographic distribution of predictive power values (Pearson’s r) of the encoding model, averaged across participants, separately for multivariate (top) and univariate (bottom) analysis. C. TRF encoding models for the first half (green) and second half (yellow) of the experiment, plotted from electrode ‘Fz’, separately for target and non-target speech. Shaded highlights denote SEM across participants. D. Speech reconstruction accuracies for the first and second half of the experiment, for both target and non-target speech. Error-bars denote SEM across participants.

The TRFs estimated separately in the 1 and 2 half of the experiment for target and non-target speech were highly similar in their spatio-temporal properties, and no significant differences between them were found (Figure 4C). An ANOVA comparing the reconstruction accuracies for target vs. non-target speech across both halves of the experiment, revealed a main effect of task-relevance [target vs. non-target: F(1,22)=28.3, p<0.001] but no main effect of half [F(1,22)=0.12, p=0.73] or interaction between the factors [half x talker: F(1,22)=0.87, p=0.87)]. These results were confirmed using a Bayesian ANOVA which indicated that the main effect of task-relevance could be embraced with high confidence and explains most of the variance (BF_inclusion_ = 317, p=0.002) but there was no reliable difference between the two halves or interaction (BF_inclusion_ = 0.26 and BF_inclusion_ = 0.27 respectively, both ps > 0.7).

Moreover, we found that decoders trained in one half of the experiment generalized well to the other half when tested on stimuli that shared the same task-relevance (role – target / non-target), but did not generalize well to stimuli that shared the same talker-identity but had different roles in the two halves of the experiment (Figure 5). Moreover, the modulation of reconstruction accuracy by task-relevance was preserved even in this cross-over analysis. This suggests that the decoders are largely invariant to talker-identity, and primarily capture features related to the role of the talker in the given task and/or their sensory properties (in this case, being presented audio-visually from a central location).

**Figure 5.**
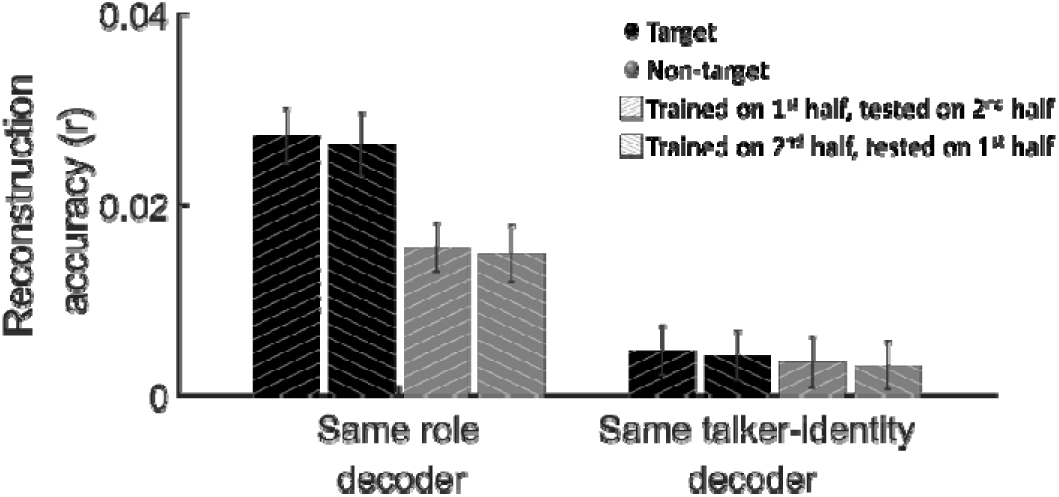
Generalizability across talkers and time. Reconstruction accuracies for decoders trained on data from one half of the experiment (either on target or non-target speech) and tested on data from the other half of the experiment, separately for same role decoders (e.g. train on target and test on target) and for same talker-identity decoders (e.g. train on male talker, test on male talker).

#### Individual-level results

Statistical analysis of speech-tracking in individual participants was three-tiered, assessing the significance of: (I) reconstruction accuracies for target and non-target speech; (II) the neural-bias index; and (III) differences between the two parts of the experiment.

Figure 6A (left panel) shows the reconstruction accuracies for target and non-target speech in individual participants, relative to their personal chance-level value (p=0.05 cutoff, derived in a data-driven manner using S-R permutation). All but one participant showed significant reconstruction accuracy of the target speech (22/23 participants - 95%), and most participants also showed higher than chance reconstruction for the non-target speech (18/23 participants - 78%). Moreover, reconstruction accuracies for target and non-target speech were positively correlated, across participants (Pearson’s r = 0.43, p = 0.038; Figure 6A right panel).

**Figure 6.**
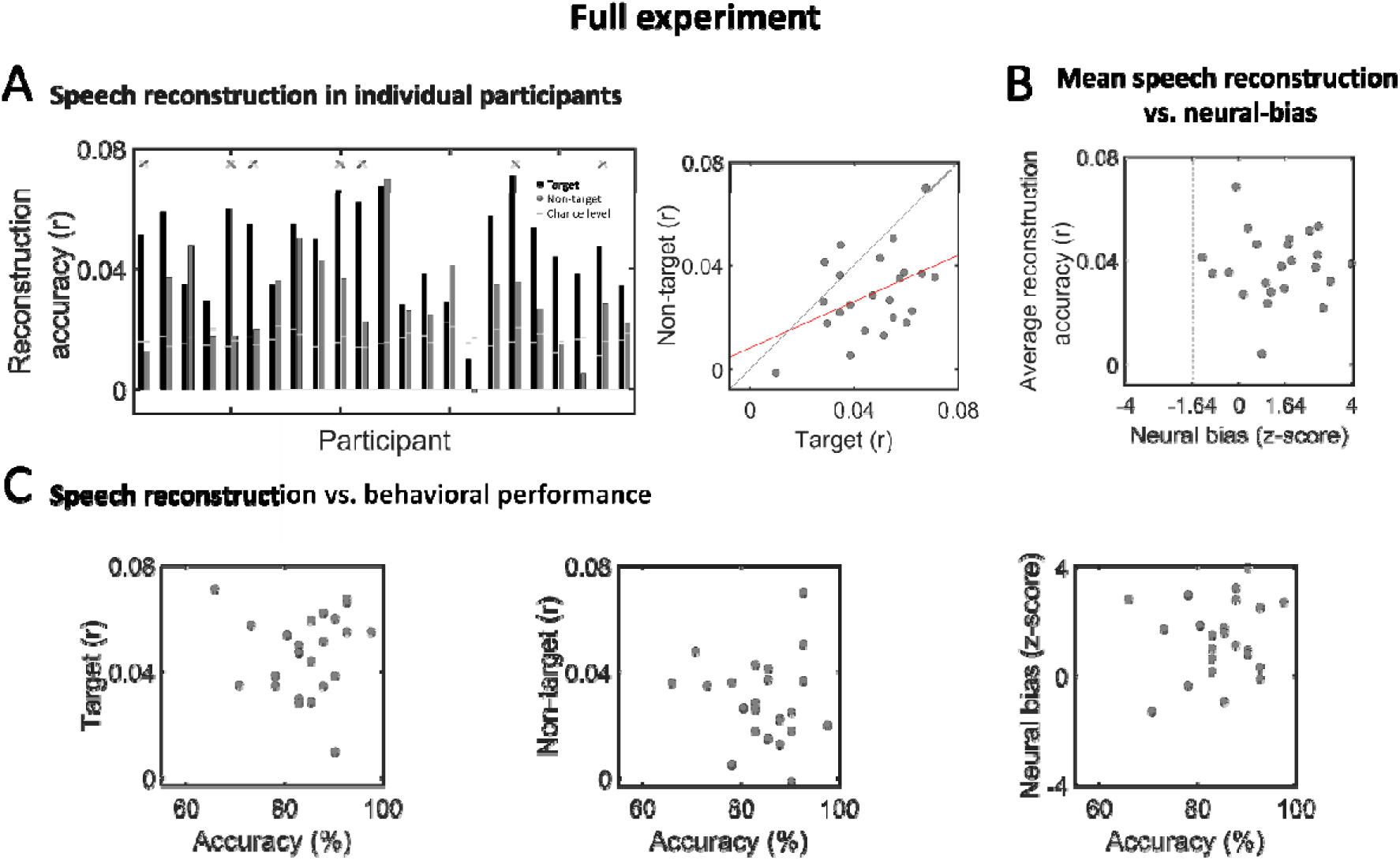
Speech Reconstruction and Neural-bias in individual participants - Full experiment. **A.** Left: Bar graphs depicting reconstruction accuracy in individual participants for target (black) and non-target (dark gray) speech. Horizontal light gray lines represent the p=0.05 chance-level, derived for each participant based on data-driven S-R permutation. Asterisks indicate participants who also showed significant neural-bias to target speech (see panel B). Right: Scatter plot showing reconstruction accuracies for target and non-target speech across all participants. The red line represents the linear regression fit between the two variables, which was significant (Pearson’s r = 0.43, p = 0.038). **B.** Scatter plot showing the average reconstruction accuracy and neural-bias index across participants, which were not significantly correlated. Vertical dashed lines indicate the threshold for significant neural-bias (z=1.64, one-tailed; p<0.05). **C.** Scatter plots showing the accuracy on behavioral task vs. reconstruction accuracy of target speech (left), non-target speech (middle) and the neural-bias index (right), across all participants. No significant correlations were found.

In Figure 6B we compare the average reconstruction accuracy from each participant to their “Neural-Bias index”, which reflects the difference in reconstruction accuracy for the target vs. non-target speech (normalized relative “target-agnostic” permutations of the data). When using a cutoff of z > 1.64 (p<0.05 one tailed), only 10/23 participants (43%) showed a significant neural-bias index, and if we use a more conservative threshold of z > 1 (p<0.15 one tailed) this proportion changes only slightly to 13/23 participants (56%). Interestingly, the reconstruction accuracies themselves and the neural-bias index were not correlated with each other (Pearson’s r=-0.017, p=0.94), suggesting that these metrics are independent.

We further tested whether performance on the behavioral tasks (answering comprehension questions) was correlated with reconstruction accuracy of either speech stimuli or the neural-bias index, however none of the brain-behavior correlations were significance (neural-bias vs. behavior: r=0.14, p=0.5; target reconstruction accuracy vs. behavior: r=0.032, p=0.88; non-target reconstruction accuracy vs. behavior: r=-0.088, p=0.69; Figure 6C).

Figures 7&8 depict the results of the same analyses shown in Figure 6, but conducted separately on data from each half of the experiment. Here, speech reconstruction was significant in a fewer proportion of participants, with 17/23 (74%) showing significant reconstruction of target speech in both the 1^st^ and 2^nd^ half (although these were not necessarily the same participants), and 10/23 (52%) or 12/23 (43%) participants showing significant reconstruction of non-target speech in the 1^st^ and 2^nd^ half of the experiment, respectively (Figures 7A & 8A). Reconstruction accuracy of target and non-target speech were not significantly correlated in the 1 half of the experiment (Pearson’s r = 0.2, p=0.33) but were in the 2 half (Pearson’s r=0.4, p=0.048).

**Figure 7.**
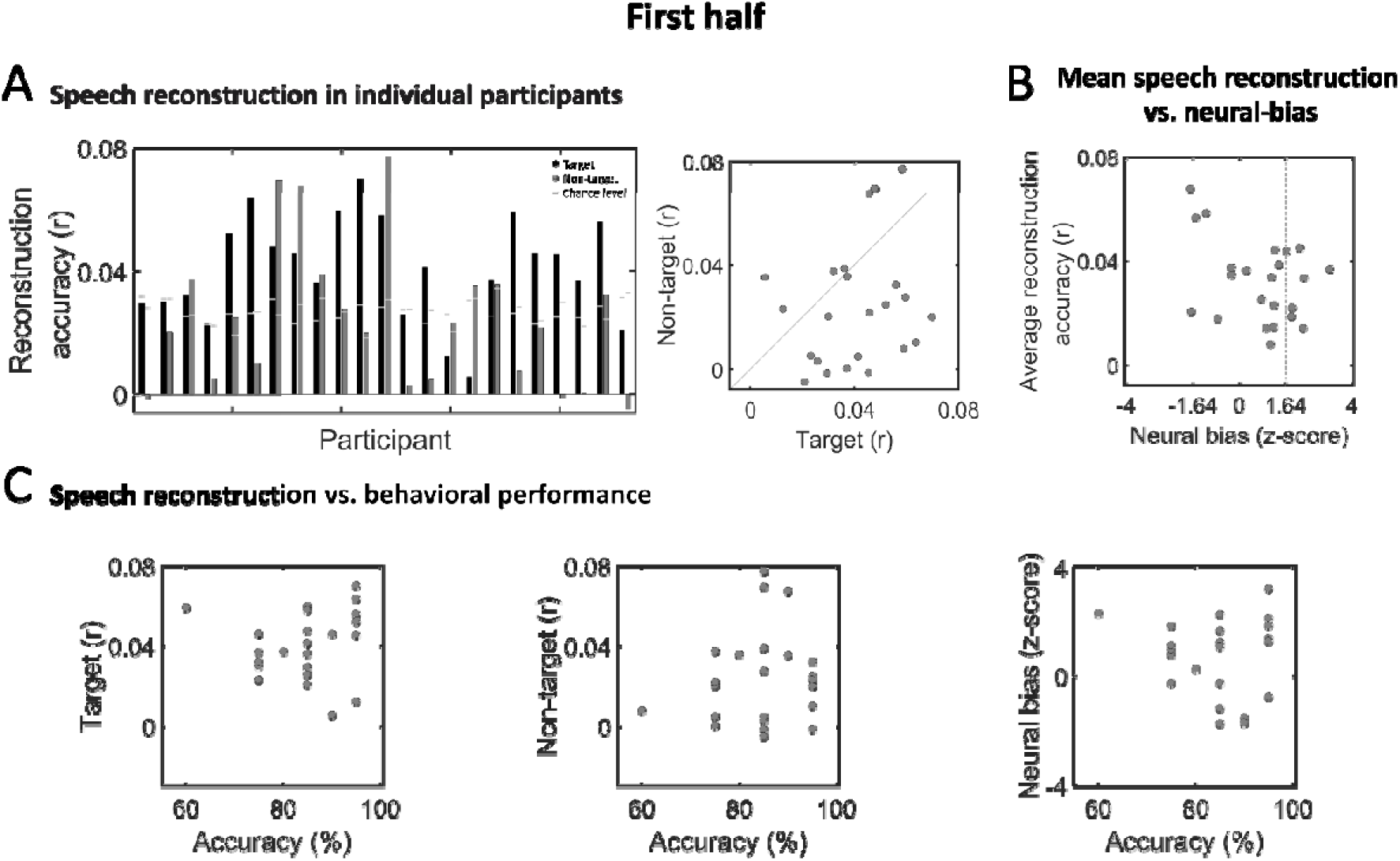
Speech Reconstruction and Neural-bias in individual participants – 1^st^ half of experiment. **A.** Left: Bar graphs depicting reconstruction accuracy in individual participants for target (black) and non-target (dark gray) speech. Horizontal light gray lines represent the p=0.05 chance-level, derived for each participant based on data-driven S-R permutation. Right: Scatter plot showing reconstruction accuracies for target and non-target speech across all participants. **B.** Scatter plot showing the average reconstruction accuracy and neural-bias index across participants, which were not significantly correlated. Vertical dashed lines indicate the threshold for significant neural-bias (z=1.64, one-tailed; p<0.05). **C.** Scatter plots showing the accuracy on behavioral task vs. reconstruction accuracy of target speech (left), non-target speech (middle) and the neural-bias index (right), across all participants. No significant correlations were found.

**Figure 8.**
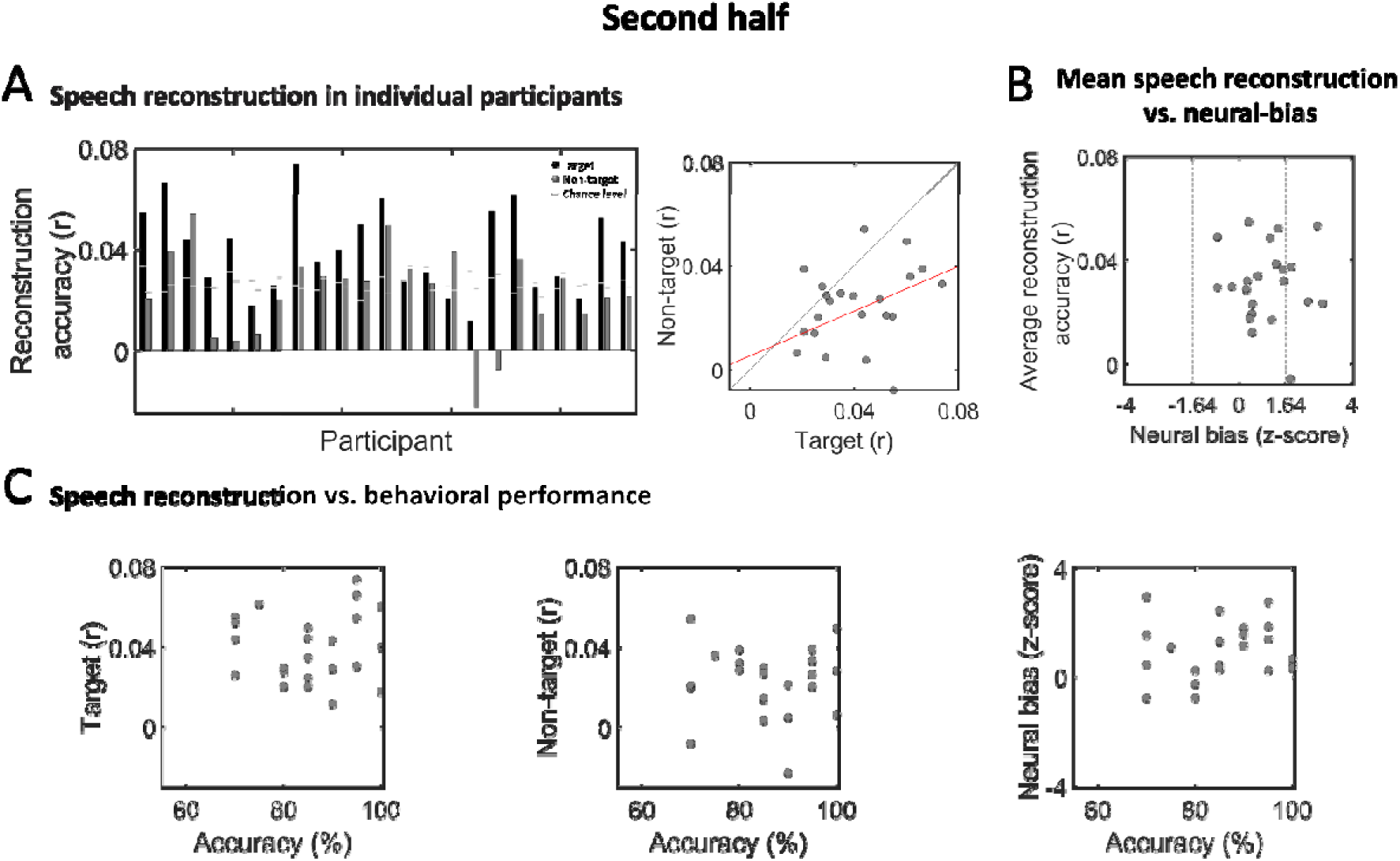
Speech Reconstruction and Neural-bias in individual participants – 2^nd^ half of experiment. **A.** Left: Bar graphs depicting reconstruction accuracy in individual participants for target (black) and non-target (dark gray) speech. Horizontal light gray lines represent the p=0.05 chance-level, derived for each participant based on data-driven S-R permutation. Right: Scatter plot showing reconstruction accuracies for target and non-target speech across all participants. The red line represents the linear regression fit between the two variables, which was significant (Pearson’s r = 0.4, p = 0.048). **B.** Scatter plot showing the average reconstruction accuracy and neural-bias index across participants, which were not significantly correlated. Vertical dashed lines indicate the threshold for significant neural-bias (z=1.64, one-tailed; p<0.05). **C.** Scatter plots showing the accuracy on behavioral task vs. reconstruction accuracy of target speech (left), non-target speech (middle) and the neural-bias index (right), across all participants. No significant correlations were found.

When evaluating the Neural-Bias Index of individual participants, we found that only 7/23 (30%) and 5/23 (21%) showed significantly better reconstruction accuracy for the target vs. non-target speech (z>1.64, p<0.05 one tailed; 1^st^ and 2^nd^ half respectively). As observed for the full experiment, here too reconstruction accuracy was not correlated with the neural-bias index in either half (1^st^ half: r=-0.37, p=0.08; 2 half: r=-0.026, p=0.9; Figures 7B & 8B), nor were any brain-behavior correlations significant (1 half: neural-bias vs. behavior: r=-0.075, p=0.73; target reconstruction accuracy vs. behavior: r=0.095, p=0.67; non-target reconstruction accuracy vs. behavior: r=0.14, p=0.54; Figure 7C; 2^nd^ half: neural-bias vs. behavior: r= 0.1, p=0.64; target reconstruction accuracy vs. behavior: r=0.05, p=0.83; non-target reconstruction accuracy vs. behavior: r= -0.007, p=0.97; Figure 8C).

Last, we compared the speech-reconstruction accuracies and neural-bias indices obtained in each half of the experiment, but found that none of these measures were significantly correlated between the 1 and 2 half of the experiment (neural-bias: Pearson’s r= -0.005, p= 0.98, target speech: Pearson’s r= 0.34, p=0.11, non-target speech: Pearson’s r=0.17, p=0.44; Figure 9). Only a handful of participants showed above-change differences between the 1 and 2 half of the experiment in these metrics, relative to a distribution of order-agnostic permuted data (shown in red in Figure 9), however these may represent false-positives due to multiple comparisons.

**Figure 9.**
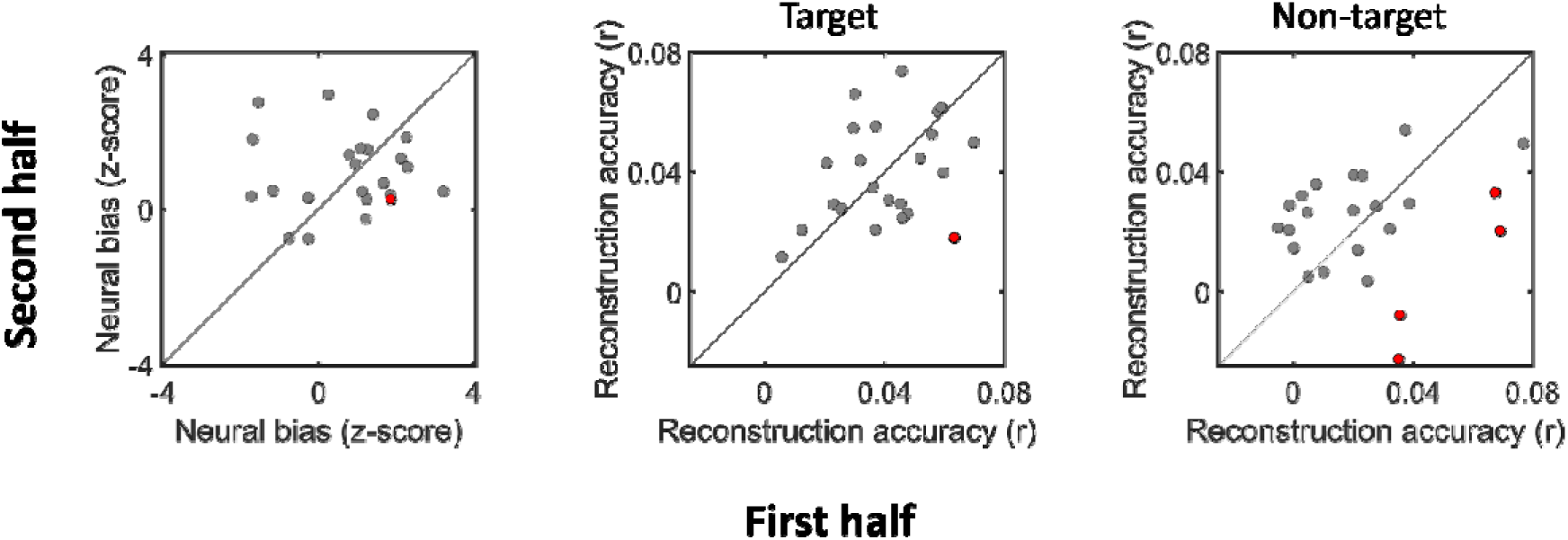
First vs second half of experiment. Scatter plots showing the neural-bias index (left), target speech (middle) and non-target speech reconstruction accuracies (right) across all participants, in the 1^st^ vs. 2^nd^ half of the experiment. No significant correlations were found for any of the measures. Participants for whom significant difference were found between the two halves (based on an order-agnostic permutation test) are marked in red.

## Discussion

Here we studied the neural representation of target and non-target speech in a spatially-realistic audiovisual setup, at both the group-level and in individual participants. The group level results are in line with results from previous two-talker experiments that used less ecological paradigms (e.g., dichotic listening, scripted speech materials etc.), namely that the acoustic envelope of target speech is represented more robustly than that of non-target speech (Brodbeck, Presacco, et al., 2018; Ding et al., 2012; Fiedler et al., 2019; Fuglsang et al., 2017; Har-shai Yahav & Zion Golumbic, 2021; Kaufman & Zion Golumbic, 2023; Kerlin et al., 2010; Mesgarani & Chang, 2012; Niesen et al., 2019; O’Sullivan et al., 2015; Straetmans et al., 2024; Zion Golumbic, Ding, et al., 2013). We also show that neural-tracking is invariant to talker identity and does not change significantly over the course of the experiment (in magnitude or spatio-temporal features of the decoder). This supports the robustness of this measure for use in free-field audiovisual contexts, laying the foundation to extend scientific investigation of selective attention to more realistic environments (Brown et al., 2023; Levy et al., 2024; Parsons, 2015; Risko et al., 2016)

However, examination of individual-level results revealed that the neural-bias observed at the group level is not seen consistently in all, or even in most, listeners. Fewer than half of the participants showed significant neural-bias for target speech, whereas in most participants non-target speech could be reconstructed just as well as target speech from the neural signal. Importantly, reconstruction accuracy and the neural-bias index were not correlated, indicating that variability across participants cannot be trivially explained by poor speech-tracking or low SNR. Moreover, neither the reconstruction accuracy of target or non-target speech nor the neural-bias index were significantly correlated with performance, suggesting they carry limited behavioral consequences. These results are similar to those reported in a previous MEG study (Kaufman & Zion Golumbic, 2023), which taken together lead us to suggests that although speech-tracking metrics are useful for studying selective attention at the group level, they may fall short as “neural-markers of selective attention” at the individual level. Below we discuss the potential implications of these results for understanding the different strategies listeners may employ to deal with concurrent speech in realistic multi-talker contexts.

### Group-level effects: highly robust and generalizable

The group-level TRFs, derived separately for target and non-target speech using an encoding approach, represent spatio-temporal filters that capture the neural responses to each of the two concurrent talkers (Ding & Simon, 2012; Fiedler et al., 2019; Fuglsang et al., 2017; Kaufman & Zion Golumbic, 2023; Power et al., 2012; Zion Golumbic, Ding, et al., 2013). They share similar fronto-central topographical distribution, which is typical of auditory responses, however, they differ in their time-course. The TRF for target speech contains three prominent peaks – roughly at 60ms, 110ms and 170ms - which is in line with previous studies and are thought to reflect a cascade of information flow from primary auditory cortex to associative and highly-order regions (Brodbeck, Presacco, et al., 2018; Chen et al., 2023). Conversely, the TRF for non-target speech showed only a single positive peak, around 70 ms, which likely reflects its early sensory encoding, but not the two later peaks. Past studies have reported mixed results regarding the temporal similarity of TRFs for target and non-target speech, some showing similar time-courses albeit with reduced amplitudes for non-target speech (Ding et al., 2012; Fiedler et al., 2019; Kaufman & Zion Golumbic, 2023; Kerlin et al., 2010; J. A. O’Sullivan et al., 2015; Zion Golumbic, Ding, et al., 2013), and others showing that the later TRF peaks are not present for non-target speech (Har-shai Yahav et al., 2023; Jaeger et al., 2020). It is likely that differences in spatio-temporal characteristics of TRFs for target and non-target speech are affected both by the specific perceptual attributes of the stimuli themselves (e.g., audiovisual vs. audio presentation, spatial location) as well as by their task-related role (target vs. non-target). In the spatially-realistic experimental design used here, these factors are inherently confounded, just as they are under real-life conditions in which listeners look at the talker that they are trying to pay attention to. A previous study by O’sullivan et al. (2019 and re-analyzed by Ahmed, Nidiffer, & Lalor, 2023) attempted to dissect the specific contribution of selective attention vs. audiovisual input by including a condition in which participants watched video of a talker that they had to ignore, and attended to a talker who they could not see. However, it is not clear to what degree such a highly artificial design (which is also extremely cognitively demanding) is representative of the mechanisms that listeners use when processing speech under natural circumstances. Instead, here, rather than trying to assert whether the differences in TRF are due to “selective attention per se” or to “perceptual differences”, we accept that in real-life these often go together. We posit that as selective attention research progresses to more ecologically valid contexts, these factors cannot (and perhaps need not!) be teased apart, but rather should be considered as making inherently joint contributions to the recorded neural signal.

The TRFs and reconstruction accuracies derived separately from each half of the experiment were highly similar, both for target and for non-target speech. This indicates that listeners were highly effective at adapting their neural encoding after the mid-way shift in the identity of the target talker and the topic of the lecture. It also demonstrates the robustness of EEG-based speech-tracking measures and of the neural-bias towards target speech when using roughly 10 minutes of data (at least at the group level). When designing this study, we postulated that neural-tracking and/or neural-bias to the target speech might be worse in the 2^nd^ half of the experiment, either due to fatigue (Jaeger et al., 2020; Moore et al., 2017) or due to higher cognitive interference of the non-target talker who had previously been the target talker (Har-shai Yahav et al., 2023; Johnsrude et al., 2013). The finding that here the switch did not carry a behavioral or neural processing cost is testimony to the high adaptability and flexibility of the attentional system, which does not ‘get stuck’ on processing previously relevant features but updates its preferences according to changing task demands (Agmon et al., 2021; Kaufman & Zion Golumbic, 2023; Kiesel et al., 2010; Koch & Lawo, 2014). This result is somewhat in contrast to some previous findings showing that speech processing was adversely affected by attention-switching. For example, a recent study similar to ours (albeit using auditory-only stimuli), found that a previously-attended stream posed more of an interference to behavior task compared to a consistently task-irrelevant stream (Orf et al., 2023). Other studies have also demonstrated reduced speech processing, decreased intelligibility, and impaired recall of specific details, resulting from attention-switching between talkers (Best et al., 2008; Getzmann et al., 2017; Lin & Carlile, 2015; Teoh & Lalor, 2019; Uhrig et al., 2022). However, in those studies the switches occurred on a per-trial basis, which likely creates more opportunities for confusion relative to the current study where the target talker was switched only once. Admittedly, there is much yet to explore regarding attention-switching between talkers, particularly under ecological conditions where switches are contextual and often initiated by the listener themselves. The current findings contribute to these efforts by demonstrating that the neural representation of target and non-target speech is invariant to talker-identity and stabilizes nicely after a switch in a realistic audiovisual context.

### From group averages to individual level responses

The vast majority of cognitive neuroscience research, and particularly when using EEG, relies on averaging results from large samples of participants to obtain group-level results. This has traditionally been motivated by the noisy-nature of many EEG metrics, the variability across individuals and the need to generalize results beyond a specific sample (Luck et al., 2000; Makeig et al., 2004; Steven J. Luck, 2005). However, increasingly, there is also a desire to derive reliable EEG-based measures from the brains of individuals - to be used, for example, to explain variability in behavioral performance and cognitive capabilities, as biomarkers for clinical symptoms, and to monitor the effectiveness of personalized interventions (Bednar & Lalor, 2020; Geirnaert et al., 2024; Hadley & Culling, 2022; O’Sullivan et al., 2017). Deriving reliable individual-level EEG-based metrics for speech processing and/or for attention has been particularly appealing, given their potential utility for clinical, educational and technological interventions. The development of speech-tracking methods over the past decade has given hope that this metric will prove to be useful for individual-level assessments. Indeed, in the domain of hearing and speech processing, several groups have demonstrated robust correlations between neural speech-tracking metrics and speech intelligibility, for example in children with dyslexia or in those with hearing impairment (e.g., Keshavarzi et al., 2022; Van Hirtum et al., 2023; Xiu et al., 2022). In contexts that require selective attention to a target talker, speech-tracking methods have also been successfully applied to distinguish between the neural representations of target and non-target speech in individual participants, and even on a per-trial basis (a classification-based approach that can be effective, for example, for designing neuro-steering devices or hearing aids; (Fallahnezhad et al., 2023; Henshaw & Ferguson, 2013; Kidd, 2017; O’Sullivan et al., 2015) (see Extended Data). However, to date fewer studies have looked at the neural-bias index in individual participants, a metric that offer insights not only into the distinction between target and non-target speech but into the modulation of their neural representations as a function of task-relevance, which is thought to be a signature of top-down selective attention (Bidet-Caulet et al., 2007; Hansen & Hillyard, 1983; Hillyard et al., 1973; Manting et al., 2020; Woods et al., 1984).

The neural-bias in reconstruction accuracies of target and non-target speech for individual participants is shown qualitatively in several previous papers, but statistical analyses focused mostly on the group-level (e.g., Ding et al., 2012; Ding & Simon, 2012; Fuglsang et al., 2017; Rosenkranz et al., 2021). Kaufman & Zion Golumbic (2023) introduced the use of attention-agnostic permutation tests to quantified and test the neural-bias index statistically in individual listeners. Results from that MEG study are similar to those found here, whereby fewer than half of the participants (∼30% in that study) exhibited significant modulation of speech-tracking by selective attention, even when using a “permissive” statistical threshold. The fact that group-level averages consistently shown a robust difference between target and non-target speech despite the underlying variability across participants, is explained by the asymmetric distribution of the neural-bias metric - since none of the participants show an ‘opposite’ bias (i.e., more accurate reconstruction of non-target vs. target speech). This asymmetric distribution also lends further credibility to the neural-bias metric as reliably capturing the relative neural representation of the two speech stimuli.

How should we interpret the variability in neural-bias across individuals? Here we ruled-out several trivial explanations, namely that variability is due to poor EEG signal quality, poor speech-tracking abilities or poor attention to the target - since we show that the reconstruction accuracies for target and non-target speech are correlated with each other, but their average is not correlated with the neural-bias index, nor is it correlated with accuracy in answering questions about the content of target speech. Instead, we offer an alternative – admittedly speculative - interpretation, that emphasizes possible variability across individuals in their de-facto allocation of processing resources among competing talkers. We know from our subjective experience that paying attention solely to one target speech and shutting out other competing stimuli can be extremely difficult. There are numerous examples that both sensory and semantic properties of non-target speech are encoded and processed, indicating that selective attention to one talker does not imply its exclusive representation (Beaman et al., 2007; Brown et al., 2023; Dupoux et al., 2003; Har-shai Yahav et al., 2023; Har-shai Yahav & Zion Golumbic, 2021; Parmentier et al., 2018; Rivenez et al., 2006; Vachon et al., 2019). Moreover, the fact that individual can divide their attention reasonably well between two concurrent speech streams if asked to do so, indicates that sufficient perceptual cognitive resources may be available to listeners to apply a multiplexed listening strategy (Agmon et al., 2021; Kaufman & Zion Golumbic, 2023; Vanthornhout et al., 2019). Given that, it is reasonable to assume that even when instructed to pay attention to only one talker, listeners may devote at least some resources to the competing non-target talker as well – either voluntarily, as in divided attention; or involuntarily (Makov et al., 2022). This notion is in line with ‘load theory of attention’, which suggests that rather than attributing attentional-selection of target stimuli/features to either ‘early’ or ‘late’ stages of processing, attention should be viewed as the dynamic allocation of available cognitive resources among competing stimuli. This allocation of resources among talkers reflects their prioritization vis-à-vis their relevance to the listener, but can also vary as a function of perceptual load, task demands and listener motivation/internal state, which we propose may underlie some of the variability observed here between participants (Gagné et al., 2017; Lavie et al., 2004; Murphy et al., 2017; Peelle, 2018; Wild et al., 2012). Along these lines, the lack of a correlation between neural-bias and performance may suggest that the perceptual and cognitive demands of the current task, which emulates those encountered in many real-life situations, left many listeners with sufficient available resources to co-represent both talkers without behavioral costs. Clearly, this interpretation is speculative and the current data are insufficient for testing its plausibility, however we offer it as a hypothesis for future studies aimed at better understanding individual differences in prioritizing between target and non-target speech under realistic circumstances and whether this variability is explained by specific personal traits, by perceptual or cognitive load or by other state-related factors (Beaman et al., 2007; Colflesh & Conway, 2007; Forster & Lavie, 2008; Lambez et al., 2020; Murphy et al., 2017; Sörqvist & Rönnberg, 2014).

Another important point to note in this regard is that speech-tracking as quantified here as the envelope-following response measured using EEG, captures only a partial neural representation of the speech, primarily reflecting encoding of its acoustic properties in auditory (Crosse et al., 2016; Ding et al., 2012; Fiedler et al., 2019; Mesgarani & Chang, 2012; Zion Golumbic, Ding, et al., 2013). Recent work that has attempted to separate between neural-tracking of acoustic- and linguistic/semantic-features of speech, has suggested that selective attention primarily affects the latter (Brodbeck, Elliot Hong, et al., 2018; Ding et al., 2018; Lachter et al., 2004), although this is not always the case (Har-shai Yahav & Zion Golumbic, 2021; Parmentier, 2008; Parmentier et al., 2018; Vachon et al., 2019). Moreover, studies that have looked at neural speech-tracking across different brain regions, using source-level MEG data or intracranial EEG (ECoG) recordings, have shown a dissociation between the sensory cortical regions that co-represent concurrent speech vs. higher order regions (e.g, anterior temporal cortex, as well as frontal and parietal regions) where attention selectivity was more prominent (Brodbeck, Elliot Hong, et al., 2018; Brodbeck, Presacco, et al., 2018; Har-shai Yahav & Zion Golumbic, 2021; Zion Golumbic, Ding, et al., 2013). Accordingly, it is possible that if we were to use a more detailed characterization of the speech stimuli, had used a more complex non-linear model or had analyzed neural responses stemming from brain regions beyond auditory cortex, we might have found more extensive evidence for neural-bias in individual participants. While this is an important limitation to consider from a basic-science perspective, the use of EEG in the current study and our focus on the acoustic envelope of speech have critical applicational value. The motivation to derive personalized neural metrics of selective attention (as opposed to group-based data) has a large practical component, such as providing tools for clinical/educational assessments and interventions. As such, these would likely involve EEG recordings (which are substantially more accessible and affordable than MEG) and – in the case of speech-processing - would rely on analyzing speech-features that are easy to derive (and do not require a tedious annotation process; Agmon et al.). The current results suggest that in order to assess the utility of the neural-bias index as a practical brain-based ‘biomarker’ of selective attention in individual listeners, we need to better understand the factors that underlie the observed variability reported here and in previous studies.

## Conclusions

In traditional cognitive-neuroscience research there is a desire to manipulate a specific construct (e.g., which stimulus is the target) while controlling for all low-level sensory differences between stimuli. However, as we turn to studying neural operation under increasingly ecological conditions, perfect control is less possible. In the case of selective attention, “targetness” is often accompanied with specific sensory attributes, making targets inherently different than non-targets. Here we studied one such case – where target speech is audiovisual and non-target speech is peripheral and auditory only. We show that under these more realistic conditions, the hallmark signature of selective attention – namely the modulation of sensory representation, and its robustness to switches in target-identity – is conserved, at least at the group level. At the same time, our results also point to an underlying diversity among participants in how that this modulation manifests, to the degree that in over half of the participants target and non-target speech were represented just as well (albeit in different ways). These results emphasize that there is still much to explore regarding how the brain – or how different brains - treats target and non-target speech when attempting to achieve selective attention. This work call for more granular investigation of how factors such as task-difficulty, perceptual load, listener motivation and personal traits ultimately affect neural encoding of competing stimuli, under ecological conditions.

