## Extended Data for "Neural Speech-Tracking During Selective Attention: A Spatially Realistic Audiovisual Study"

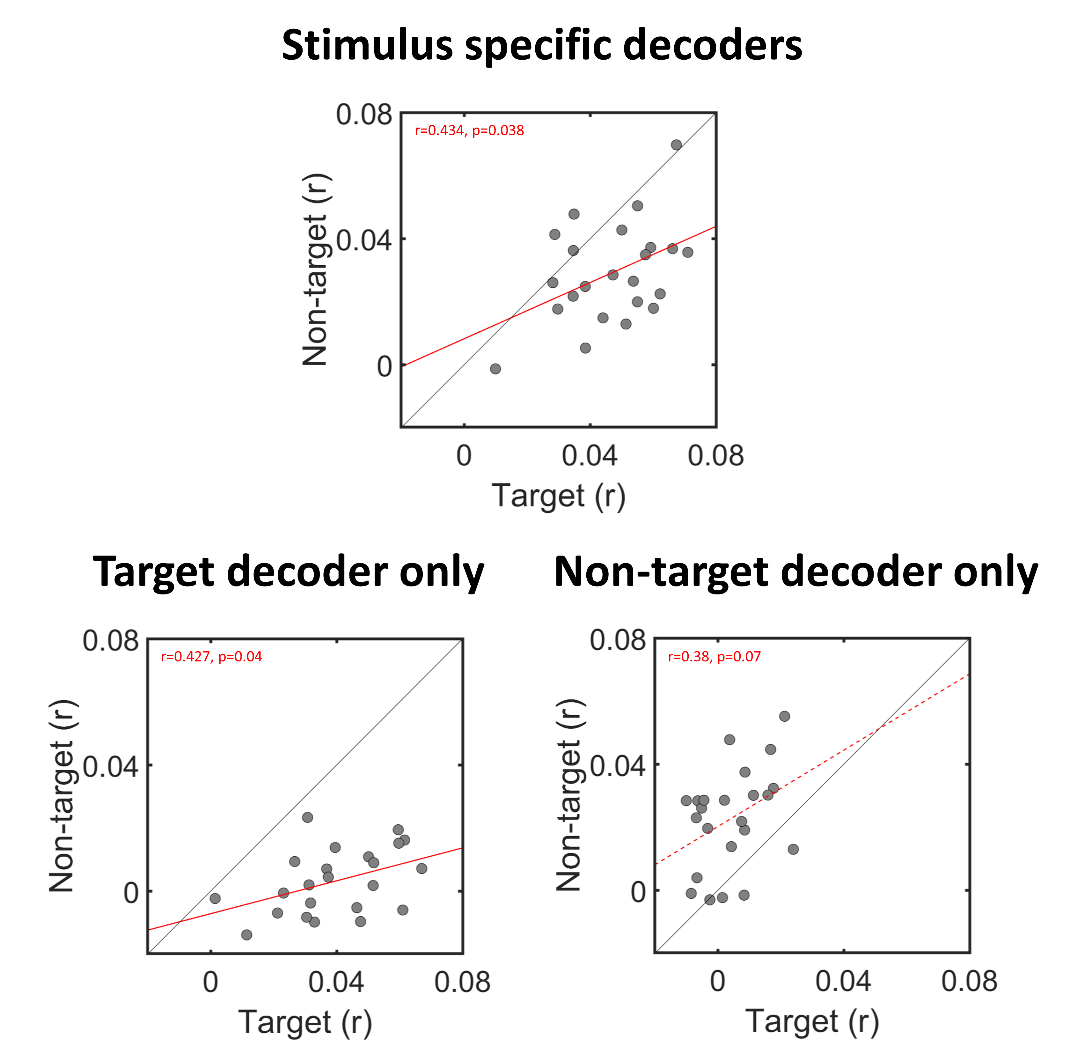

**Figure S1.** **Comparison of** **decoder-testing approaches.** Here we compare two approaches for testing the performance of decoders trained on EEG data to reconstruct the envelope of concurrently presented speech.

***Top***: The approach used and reported in the current study, in which two **Stimulus specific decoders** were trained using a multivariate approach to reconstruct the envelopes of target and non-target speech presented concurrently. The scatter-plot shows reconstruction accuracies achieved for both decoders across all participants, when tested on left-out data of the same type (i.e., how well the target decoder can reconstruct left-out target speech, and how well the non-target decoder can reconstruct left-out non-target speech). The gray line reflects the diagonal, and the red line represents the linear regression fit between the two variables which was statistically significant [data is the same as in Figure 6A].

***Bottom***: Re-analysis of the same data using the auditory attention-decoding (AAD) approach, in which a decoder is trained only on one stimulus (e.g., on target speech), and is then tested on left-out data of the same stimulus (target) and of the other stimulus (non-target), and the two results are compared for classification purposes. The left panel shows a scatter-plot showing how well a decoder trained on target speech can reconstruct left-out target speech vs. how well it can reconstruct left-out non-target speech, across all participants. The right panel shows the same for a decoder trained on non-target speech. The gray line reflects the diagonal, and the red line represents the linear regression fit between the two variables (dashed line indicates a marginally significant regression).

In this analysis, almost all dots fall either below or above the diagonal, clearly showing between reconstruction performance when a decoder is tested on data of the same type that it was trained on. This is in line with multiple studies, that propose using this approach for practical applications, such as controlling a neuro-steered hearing device (Fallahnezhad et al., 2023; Henshaw & Ferguson, 2013; Kidd, 2017; J. A. O’Sullivan et al., 2015).

Given the qualitative sensory differences between target and non-target speech in the spatially-realistic audiovisual setup used (e.g., spatial location, audio/audio-visual presentation etc.), it is not very surprising that decoders trained on these stimuli would differ from each other. However, we posit that this approach is less appropriate in the current study, where the goal was not just to distinguish between the two stimuli, but to test whether target speech is represented more robustly in the neural data than non-target speech, a pattern that is considered a signature of ‘selective attention’ - i.e., enhancement of target speech and/or suppression of non-target speech (Ding et al., 2012; Fiedler et al., 2019; Kerlin et al., 2010; J. A. O’Sullivan et al., 2015; Zion Golumbic, Ding, et al., 2013). For this purpose, we believe that is it more appropriate to optimize decoders for each stimulus separately (thus accounting for their differences in properties), and then assess how well each one performs for predicting the stimulus it was trained on (the model’s goodness-of-fit/predictive power/accuracy). Using this approach, if we find that both decoders perform very well – this indicates that both stimuli are represented with similar precision in the neural response. Conversely, finding that the decoder for one stimulus outperforms the other can be interpreted as superior or more detailed neural encoding of that stimulus relative to the other, effects that have been associated with better intelligibly and/or higher levels of attention to this stimulus (Best et al., 2008; Getzmann et al., 2017; Lin & Carlile, 2015; Orf et al., 2023; Teoh & Lalor, 2019; Uhrig et al., 2022) .
